# AI-guided discovery of antimicrobial peptides for urinary tract infections leveraging a new catalogue of the human urinary microbiome

**DOI:** 10.64898/2026.08.05.741749

**Authors:** Shanlin Ke, Franz Georg Zingl, Xu-Wen Wang, Vanessa L. Hale, Scott T. Weiss, Matthew K Waldor, Yang-Yu Liu

**Author notes:** These authors contributed equally (Shanlin Ke, Franz Georg Zingl). Correspondence should be addressed to: S.K. and Y.-Y.L.

## Abstract

Urinary tract infections (UTIs) are common infections that pose a critical burden on healthcare and society. Despite growing recognition that the human urinary tract harbors its own microbiome, its composition, functional potential, and alterations in UTI remain limited. Here, we leveraged the publicly available whole-metagenome shotgun sequencing data from 450 urinary microbiome samples collected in four independent cohorts together with genome assembly and metagenomic binning to construct an extensive human urinary microbiome catalog consisting of ∼1.3 million non-redundant microbial genes and 705 non-redundant metagenome-assembled genomes (nrMAGs). We found that microbiomes from patients with UTI carry significantly more genes linked to antibiotic resistance and virulence vs controls. There was an enrichment of multiple *Escherichia* strains in patients with UTI from two independent case-control cohorts. UTIs are becoming multidrug-resistant, and we used machine learning models to identify potential antimicrobial peptides (AMPs) in 705 nrMAGs. Furthermore, we experimentally demonstrated that two of these AMPs exhibited strong inhibitory activity against uropathogenic *Escherichia coli* strains. Our study provides a valuable resource for studying the human urinary microbiome and suggests urinary microbiome-derived AMPs represent a source of new therapeutics for UTIs.

## INTRODUCTION

Human microbiome research has predominantly focused on the gut microbiome^1^, with other body sites, such as the urinary tract, receiving comparatively limited attention^2^. The existence of a urinary microbiome has been controversial because of the long-standing dogma that “healthy” urine is sterile^3^. However, recent research has revealed that the urinary tract of healthy individuals harbors a microbial community or microbiome^2,4,5^. For example, a study demonstrated the presence of a “core” urinary microbiome in voided midstream urine samples from a population of healthy individuals ranging from 26 to 90 years^6^. These findings have sparked great interest in understanding the composition and function of the urinary microbiome and its potential implications for urinary tract health.

Urinary tract infections (UTIs) are common microbial infections, accounting for about 40% of all hospital-acquired infections^7,8^. As a significant public health concern, UTIs impact over 400 million individuals annually worldwide^9^ and impose a substantial clinical and economic burden^10^. Moreover, it is estimated that approximately 20-40% of women who experience a UTI will have an additional episode, 25-50% of whom will experience multiple recurrent episodes^11^. Increasing evidence suggests that the urinary microbiome may influence susceptibility to UTIs, disease progression, and treatment outcomes^12,13^. However, how urinary microbial communities interact with their hosts and contribute to UTIs remain poorly understood.

Current UTI diagnosis relies primarily on a combination of clinical symptoms and a positive urinary analysis or culture^14^. Antibiotics remain the standard treatment, with culture-based antimicrobial susceptibility testing guiding therapeutic decisions^15^. However, the increasing prevalence of antimicrobial resistance (AMR) among urinary pathogens poses a growing challenge to effective UTI management^16,17^. For example, uropathogenic *Escherichia coli* (UPEC), the predominant cause of UTIs, has shown increasing resistance to commonly prescribed antibiotics, including fluoroquinolones, trimethoprim-sulfamethoxazole, and cephalosporins^18^. These challenges underscore the urgent need to better understand urinary microbial ecosystems and identify alternative therapeutic strategies beyond conventional antibiotics.

Achieving this goal requires comprehensive, culture-independent characterization of urinary microbial communities^19^, as conventional urine culture primarily detects readily cultivable microorganisms while overlooking fastidious and uncultured taxa^12^. Advances in high-throughput sequencing technologies, such as 16S rRNA gene sequencing^20–22^ and whole metagenome shotgun (WMS) sequencing^2,23,24^, provide efficient and culture-independent tools to study the human urinary microbiome. However, 16S rRNA gene sequencing lacks sufficient taxonomic resolution at the species and strain levels^25^. While the WMS sequencing approach offers greater resolution, previous studies have mainly relied on reference-based methods^26,27^, whose accuracy depends on the breadth and depth of the reference databases. Importantly, the human urinary microbiome may be underrepresented in current reference databases, primarily due to its relatively limited exploration, posing a potential challenge for human urinary microbiome research.

A method to circumvent limitations imposed by reliance on reference databases is coupling *de novo* assembly of shotgun metagenomic reads with their binning into metagenome-assembled genomes (MAGs). This approach represents a culture-independent method that can efficiently identify microorganisms that have yet to be isolated and cultured and hence are absent from the current reference genome databases. Recent studies have reconstructed over 150,000 and 72,000 high-quality bacterial genomes from gut^28^ and oral^29^ microbiome samples, respectively, yielding new insights into bacterial function and prospects for future investigations. Importantly, the reconstructed MAGs provide direct knowledge of genomic contents and offer a unique opportunity to study the biosynthetic potential of microbial communities^30^, including the discovery of bioactive molecules such as antimicrobial peptides (AMPs) that may serve as novel therapeutic candidates.

In this study, we used metagenome assembly and binning strategies to reconstruct microbial genomes directly from DNA sequences of human urinary microbiome samples. Our goals were to construct an integrated gene and genome catalog to understand the human urinary microbiome, identify microbial features related to UTIs, and discover novel AMPs that could be of benefit for treatment of UTIs (**Fig. 1**). To achieve these goals, we constructed an extensive human urinary microbiome catalog (HUMC) consisting of 1,294,428 non-redundant microbial genes and 705 non-redundant MAGs (nrMAGs) at the strain level. HUMC enables us to characterize the taxonomic and functional profiles of the human urinary microbiome and identify UTI-related microbial features. Moreover, we leveraged deep learning models to identify candidate AMPs from the MAGs in the HUMC. Finally, we demonstrated that two candidate AMPs can indeed inhibit the growth of uropathogenic *E. coli* strains. These findings greatly expand our understanding of the genomics of the human urinary microbiome and its potential contributions to UTIs; moreover, they underscore its potential promise as a resource for developing therapeutic and preventive strategies for combating UTIs.

**Figure 1.**
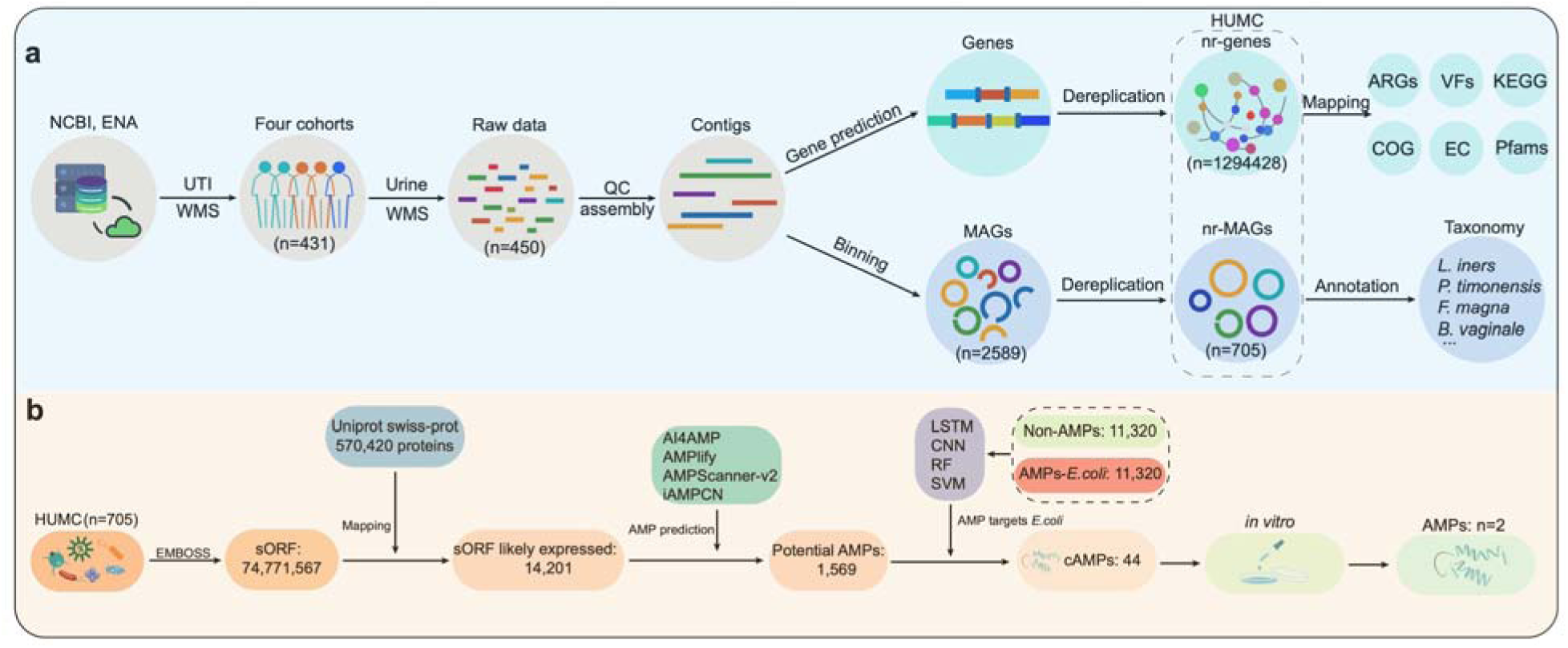
Diagram illustration of the study workflow. **a**, Construction of the human urinary microbiome catalog (HUMC). Publicly available urinary metagenomic sequence data from the National Center for Biotechnology (NCBI) and the European Nucleotide Archive (ENA) were first collected. The raw sequencing data underwent quality control and assembly, after which the contigs were used to construct the HUMC, which contains both a gene catalog and a genome catalog. Functional annotation of the genes was performed using multiple databases. Taxonomy annotation of nrMAGs was based on GTDB database. **b**. Computational framework to identify potential antimicrobial peptides (AMPs) in the HUMC. Candidate AMPs were further filtered using multiple machine learning models (acronyms), narrowing them down to a select few for chemical synthesis and experimental testing. Finally, the top candidates were tested against uropathogenic *E. coli* strains and other common pathogens to assess their antimicrobial activity. UniProt Swiss-Prot: curated protein sequence database within UniProt; KEGG: Kyoto Encyclopedia of Genes and Genomes; COG: clusters of orthologous groups; EC: enzyme commission (enzyme classification system); Pfam: protein families database; WMS: whole metagenome sequencing; QC: quality control; HUMC: human urinary microbiome catalog; nr-genes: non-redundant genes; nr-MAGs: non-redundant metagenome-assembled genomes; MAGs: metagenome-assembled genomes; sORF: small open reading frame; ARGs: antibiotic resistance genes; VFs: virulence factors; AMPs: antimicrobial peptides; cAMPs: candidate antimicrobial peptides; EMBOSS: European Molecular Biology Open Software Suite; LSTM: long short-term memory; CNN: convolutional neural network; RF: random forest; SVM: support vector machine.

## RESULTS

### Summary of human urinary metagenomic datasets

We first retrieved all microbiome studies that assessed the microbial composition of urinary samples through WMS sequencing (with data publicly available as of December 2022). The WMS sequencing data from a total of 450 urinary metagenomes from four independent cohorts were collected (Moustafa^23^, Adu-Oppong^31^, Adebayo^2^, and Neguent^32^). The data included non-UTI controls, people with a history of recurrent UTIs, and patients with active UTI (**Fig. 2a**).

**Figure 2.**
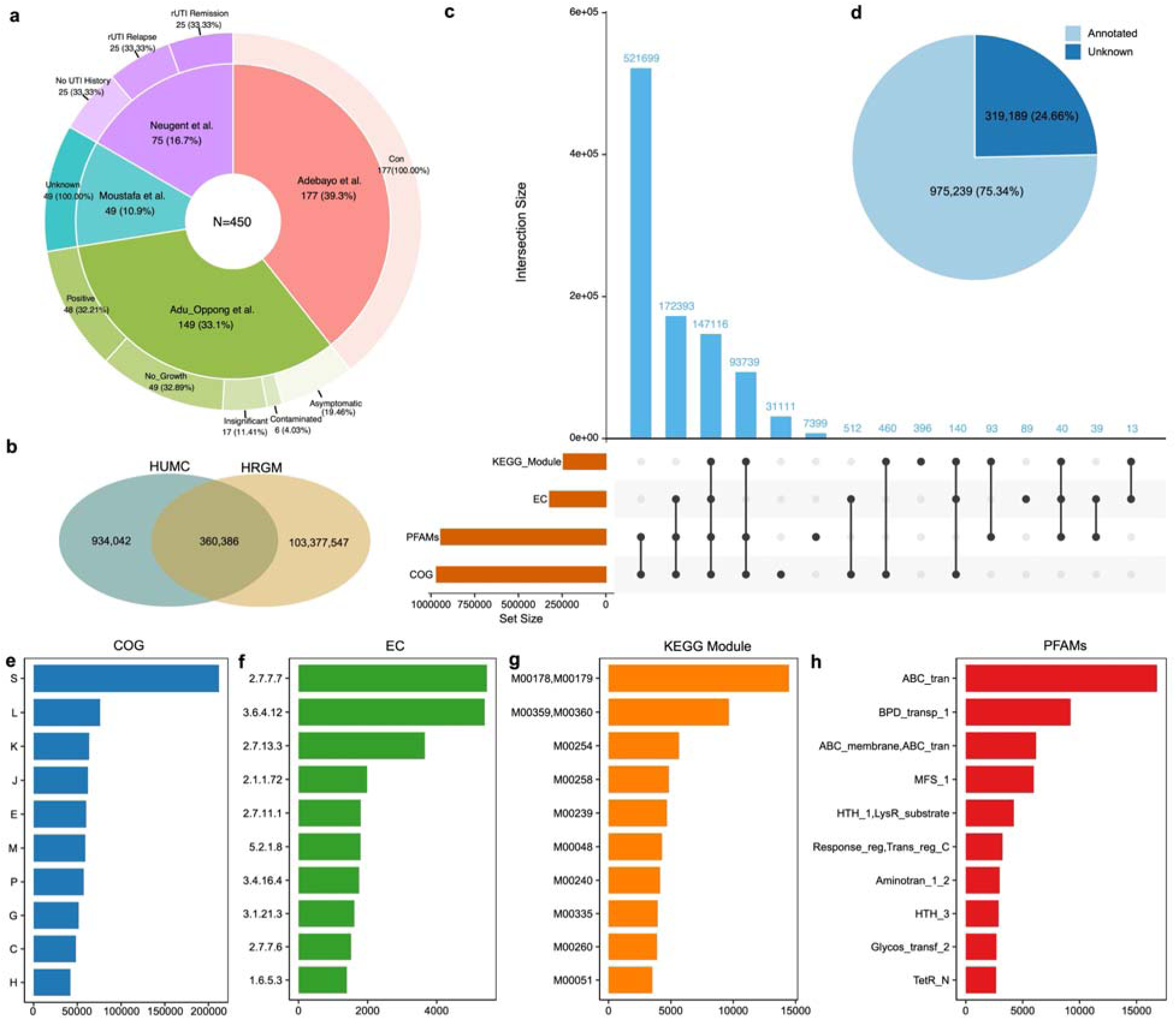
Study cohort and functional annotation of gene catalog. **a**. Sample distribution among different datasets and disease status. **b.** Comparison between genes from the HUMC and genes from the human gut reference genome (HRGM or HGRM) catalog. **c.** Number of proteins with functional annotation across five database and their degree of overlap. **d**. Number of genes being annotated. **e**. Top 10 protein clusters annotated by Clusters of Orthologous Genes (COG) database. S: functional unknown; L: replication, recombination and repair; K: transcription; J: translation, ribosomal structure and biogenesis; E: amino acid transport and metabolism; M: cell wall/membrane/envelope biogenesis; P: inorganic ion transport and metabolism; G: carbohydrate transport and metabolism; C: energy production and conversion; and H: coenzyme transport and metabolism. **f**. Top 10 Enzyme Commission classes. EC2.7.7.7: DNA-directed DNA polymerase; EC3.6.4.12: DNA helicase; EC2.7.13.3: histidine kinase; EC2.1.1.72: site-specific DNA-methyltransferase (adenine-specific); EC2.7.11.1: non-specific serine/threonine protein kinase; EC5.2.1.8: peptidylprolyl isomerase; EC3.4.16.4: serine-type D-Ala-D-Ala carboxypeptidase; EC3.1.21.3: type I site-specific deoxyribonuclease; EC2.7.7.6: DNA-directed RNA polymerase; and EC1.6.5.3: NADH:ubiquinone reductase. **g**. Top 10 KEGG modules. M00178,M00179: Ribosome; M00359,M00360: aminoacyl-tRNA synthesis; M00254: ABC-2 type transport system; M00258: Putative ABC transport system; M00239: Peptides/nickel transport system; M00048: De novo purine biosynthesis, PRPP + glutamine => IMP; M00240: Pyrimidine metabolism; M00335: secretion (sec) system; M00260: DNA polymerase III complex, bacteria; and M00051: De novo pyrimidine biosynthesis, glutamine (+ PRPP) => UMP.

### Identification of 1,294,428 non-redundant microbial genes

After quality control of the WMS sequencing data, we conducted metagenomic assembly and gene prediction to identify urinary microbial genes^33,34^. After the filtration of incomplete genes, 2,829,729 complete genes were identified across all urinary microbiome samples. These genes were then clustered at 100% amino acid identity, resulting in a non-redundant HUMC gene catalog containing 1,294,428 genes. To understand how the urinary microbiome differs from the human gut microbiome at the gene level, the microbial genes from the human reference gut microbiome (HRGM)^35^, which contains 103,737,933 genes were compared to the urinary gene compendium. Interestingly, only 27.84% (360,386/1,294,428) of genes from the HUMC were shared with the HRGM (**Fig. 2b**).

Next, the non-redundant genes were functionally annotated using eggNOG-mapper^36^. The COGs (Clusters of Orthologous Genes), KEGG (Kyoto Encyclopedia of Genes and Genomes) module, PFAMs (protein families), and ECs (Enzyme Commission) annotations were derived from the eggNOG-mapper results. Overall, 975,239 non-redundant genes from the HUMC (75.34%) could be matched to at least one of the four functional databases, whereas the remaining 319,189 genes (∼25% of the total) could not be matched, suggesting the HUMC contains a substantial number of genes whose functions remain to be determined. (**Fig. 2c-d**). Consistent with this observation, the largest COG in the HUMC was ‘gene of unknown function (**Fig. 2e**). As expected, a large proportion of genes with a known COG function present in HUMC were involved in general house-keeping functions, including replication/recombination/repair, transcription, and translation/ribosome structure and biogenesis (**Fig. 2e**). The most abundant ECs included DNA-directed DNA polymerase (EC: 2.7.7.7) and DNA helicase (EC: 3.6.4.12) (**Fig. 2f**). Among the KEGG modules, the most prevalent were M00178, M00179 (Ribosome, bacteria/archaea) and M00359, M00360 (aminoacyl-tRNA synthases, prokaryotes) (**Fig. 2g**). ABC transporters (ABC_tran) and Binding-protein-dependent transport system inner membrane component (BPD_transp_1) were the predominant protein families (**Fig. 2h**).

### Patients with UTI carry more antimicrobial-resistant genes and virulence factors in their urinary microbiome

AMR is one of the top global public health threats^37–39^. Multiple studies have reported a high prevalence of AMR among bacterial isolates from the urinary tract^40,41^. To assess AMR in the human urinary microbiome, we annotated the genes in HUMC using the Comprehensive Antibiotic Resistance Database (CARD)^42^. A total of 5,321 genes mapped to CARD, encompassing 433 antimicrobial-resistant genes (ARGs). Among these genes, the most common ARGs were associated with resistance to aminoglycoside, tetracycline, glycopeptide, and fluoroquinolone antibiotics.

We next calculated the prevalence of ARGs in two cohorts with well-defined case-control settings (i.e., Adu-Oppong^31^ and Neguent^32^, see **Methods**). Among the top 10 ARGs with varying prevalence, all exhibited higher prevalence in UTI samples, such as *Ecol_mdfA*, *emrR*, and *LptD* (**Fig. S1**). We further identified 43 and 97 differentially abundant ARGs from the Adu-Oppong and Neugent cohorts, respectively (**Fig. 3a**). Specifically, 42 differentially abundant ARGs were shared between the two cohorts. All 42 ARGs showed significantly higher abundance in UTI samples compared to control samples in both cohorts. These differentially abundant ARGs also covered those ARGs with varying prevalence, such as *cpxA*, *mdtA*, *LptD*, *mdtN*, *emrR*, and *emrA*. These findings likely reflect the repeated exposure of patients with UTI to antibiotics and suggest that novel preventative and therapeutic strategies targeting the urinary microbiome may provide beneficial approaches for UTI management.

**Figure 3.**
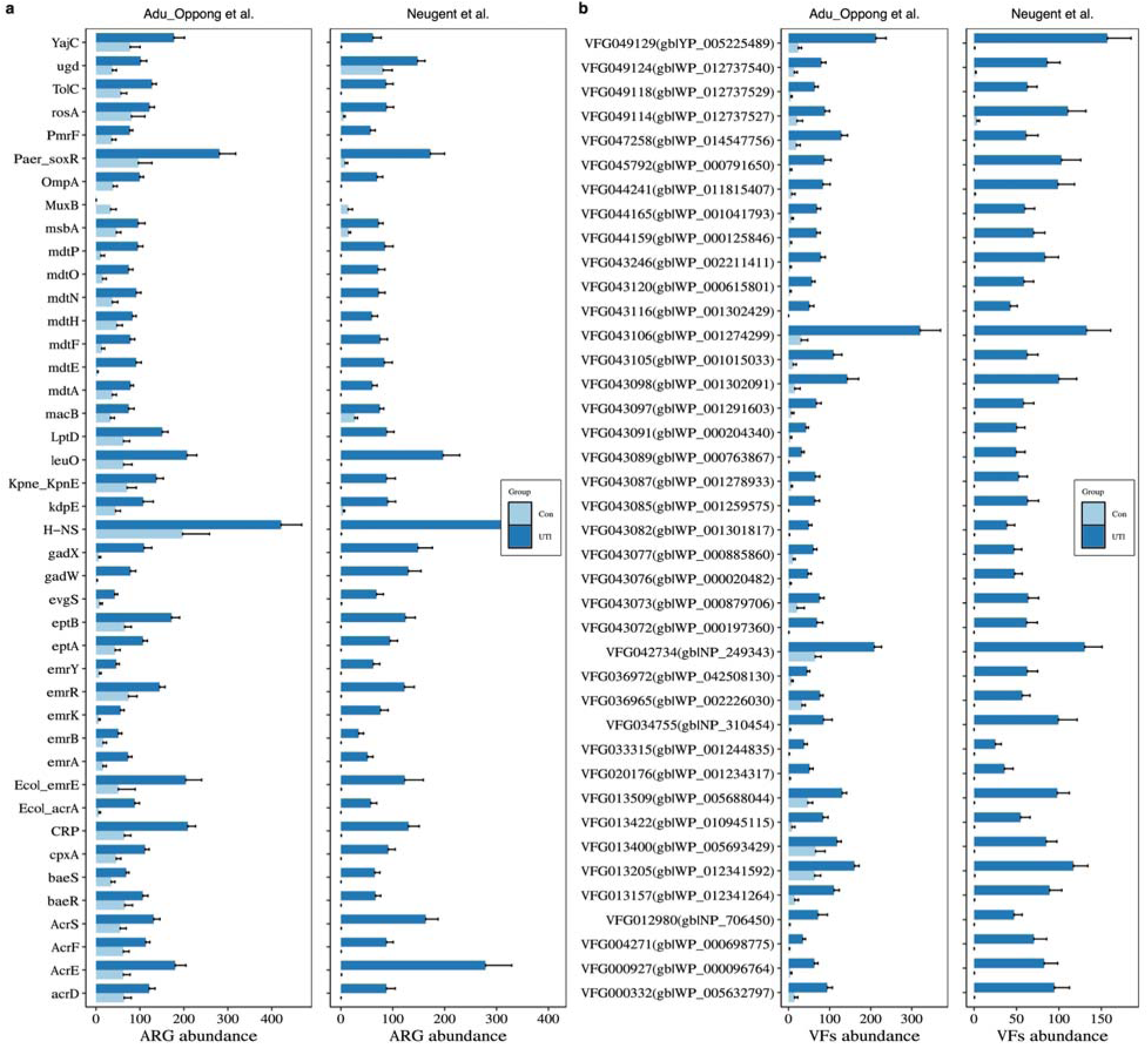
Patients with UTI carried more ARGs and VFs in their urinary microbiome compared to non-UTI individuals. **a**. Abundance (TPM) of 42 overlapped differential abundant ARGs identified from Adu-oppong and Neugent cohorts. **b**. Abundance of top 40 (based on adjusted *P* value) overlapped differential abundant ARGs identified from Adu-oppong and Neugent cohorts. Data are presented as meanO±Ostandard error of mean. P-values were calculated by two-sided Wilcoxon–Mann–Whitney test with Benjamini–Hochberg correction. All adjusted *P*-values were less than 0.01.

Virulence factors assist bacteria in colonizing their hosts and contribute to their pathogenicity^43,44^. We functionally annotated the genes from the HUMC with virulence factors in the Virulence Factor Database (VFDB)^45^. A total of 31,931 genes mapped to the VFDB, corresponding to 4,991 virulence factor genes (VFGs). The primary functional categories encoded by these VFGs were as follows: adherence (20.00%), effector delivery system (19.35%), nutritional/metabolic factors (18.01%), and immune modulation (16.93%). Similar to the observations from ARG analysis, the top 10 VFGs exhibited differential prevalence between UTI and control samples in both the Adu-Oppong and Neugent cohorts (**Fig. S2**). We further identified 272 and 487 differentially abundant VFGs from the Adu-Oppong and Neugent cohorts, respectively (**Fig. 3b**). In particular, 211 shared VFGs were identified across the two cohorts. Importantly, the top 40 VFGs were all enriched in UTI samples, consistent with the expected enrichment of urinary pathogens in those samples.

### Construction of 705 microbial genomes from human urinary samples

To deepen understanding of the microbial community residing in the human urinary tract, we employed a combination of metagenomic assembly and binning on the WMS sequencing data of 450 samples from the four cohorts. After refinement, we constructed 2,589 MAGs from these samples. These MAGs were then dereplicated into 705 non-redundant MAGs (nrMAGs) at an Average Nucleotide Identity (ANI) threshold of 99% (**Fig. 4a-b**). Among these 705 strain-level nrMAGs, 262 (37.16%) met the medium-quality criteria (50% ≤ completeness < 90% and contamination≤5%), and 443 (62.84%) were high-quality (completeness ≥ 90% and contamination ≤ 5%) (**Table S1**). The average characteristics of the nrMAGs in our dataset included a genome size of 1.83 Mb (**Fig. 4c**), 1,769.60 contigs (**Fig. 4d**), an N50 value of 93.72 Kbp (**Fig. 4e**), and a GC content of 36.68% (**Fig. 4f**). According to the Genome Taxonomy Database (GTDB)^46^, 12 phyla, 95 genera, and 200 species were identified (**Fig. 4g**). The dominant phyla were Actinobacteriota, Firmicutes (Bacillota), Bacteroidota, and Proteobacteria (**Fig. 4a**). Notably, 83 nrMAGs lacked species-level reference genomes in the GTDB (**Fig. 4h**). The species with the highest strain richness (i.e., the number of associated nrMAGs) were *Lactobacillus iners*, *Prevotella timonensis*, *Peptoniphilus_A harei_A*, *Finegoldia magna_H*, and *Bifidobacterium vaginale_G* (**Fig. 4i**).

**Figure 4.**
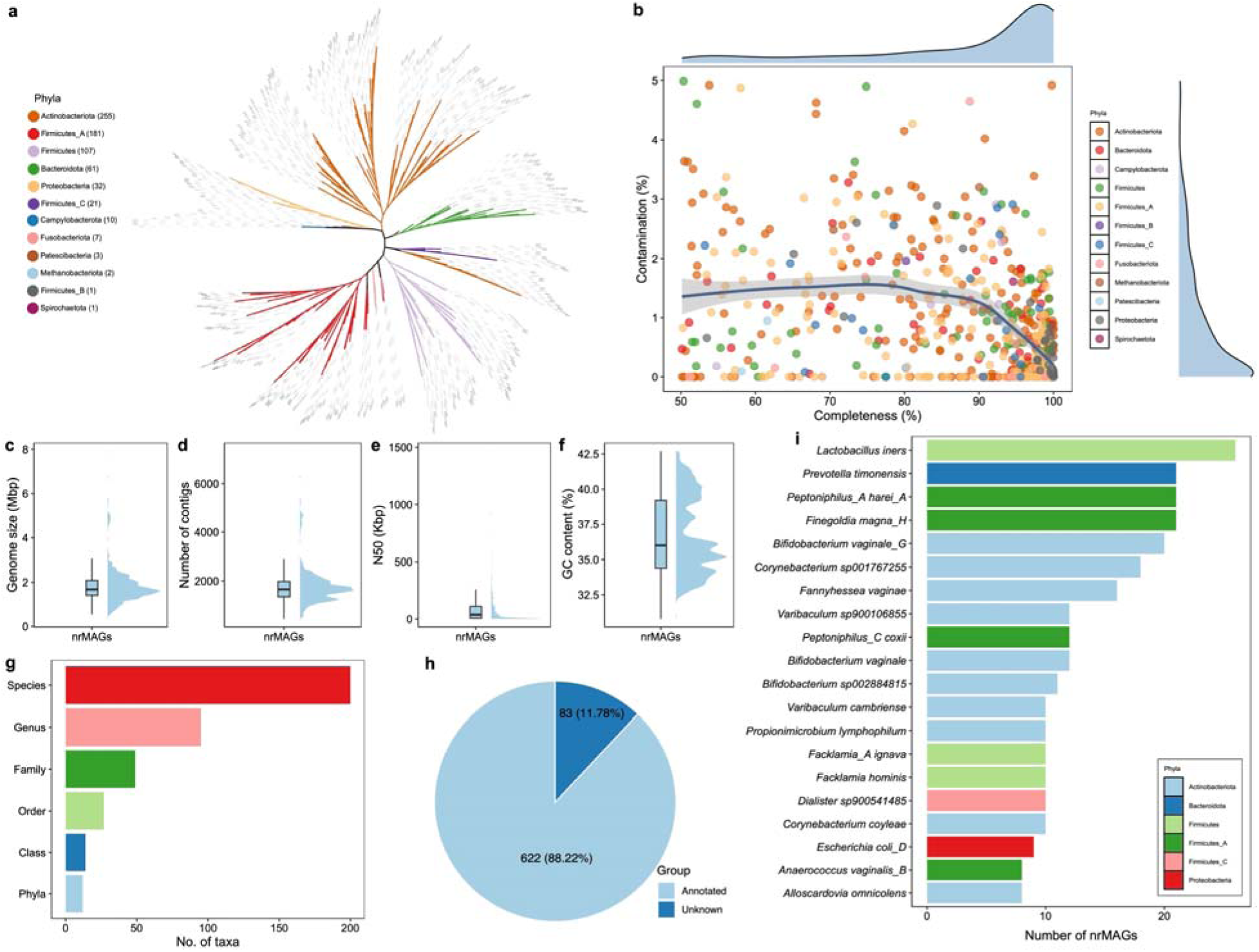
Properties of 705 high-quality non-redundant MAGs reconstructed from the human urinary microbiome samples. **a.** Phylogenetic tree of nrMAGs. The color of clades represents phylum. **b.** The distribution of completeness and contamination in nrMAGs and the color of point represents phylum. Genomic features of genomes on size (**c**), number of contigs (**d**), N50 (**e**), GC content % (**f**). **g**. Taxonomy information of nrMAGs. **h**. Number of nrMAGs with species annotation. **i**. The top 20 species with the highest strain-richness (i.e., number of nrMAGs).

### The human urinary microbiome in non-UTI samples

Consistent with previous findings that the human urinary tract of non-UTI individuals hosts an indigenous microbiome^2,47^, we identified a variable number of nrMAGs (or strains) in non-UTI samples from three cohorts (Adu-Oppong^31^, Adebayo^2^, and Neguent^32^) (**Fig. S3)**. Mapping the nrMAGs across taxonomic levels revealed Enterobacteriaceae, Staphylococcaceae, Lactobacillaceae, Bifidobacteriaceae, and Flavobacteriaceae as the dominant families (**Fig. S4**). At the genus level, the most abundant genera were *Staphylococcus*, *Lactobacillus*, *Klebsiella*, *Escherichia*, and *Bifidobacterium* (**Fig. S5**). The dominant species included *Staphylococcus aureus*, *Klebsiella pneumoniae*, *E. coli*, *B. vaginale*, *L. iners*, *L. paragasseri*, and *L. crispatus* (**Fig. S6).** Calculation of the prevalence of nrMAGs in non-UTI samples from different cohorts (**Fig. S7**), showed that some nrMAGs from *L. iners*, *E. coli*, *Streptococcus anginosus*, and *B. vaginale_G* were prevalent in non-UTI individuals. For example, across the Adu-Oppong, Adebayo, and Neguent cohorts, *L. iners* was detected in 31.03%, 11.86%, and 28.00% of samples, respectively, whereas *E. coli* was detected in 48.28%, 28.25%, and 32.00% of samples, respectively. These prevalence patterns are broadly consistent with prior reports^2^, which similarly identified these taxa as recurrent members of the urinary microbiome in non-UTI individuals.

### Identification of human urinary microbes associated with UTIs

To identify urinary microbial signatures associated with UTIs, we compared nrMAG abundance between UTI cases and controls in study cohorts with well-defined case-control cohorts. Specifically, we focused on the Adu-Oppong and Neugent cohorts, which included clearly defined UTI and control groups. In the Adu-Oppong cohort, only symptomatic patients confirmed by *in vitro* culture (growth of 1–2 uropathogenic species at ≥10^5^ colony forming units per mL) were included in the case-control comparison. In the Neugent cohort, individuals were categorized into three groups: control (no UTI history), remission (recent history of recurrent UTI but no active UTI at the time of urinary sample collection), and relapse (history of recurrent UTI and an active, symptomatic UTI at the time of urinary sample collection). The differentially abundant nrMAGs were then identified using ANCOM^48^ across the four comparisons from the two cohorts, with each cohort analyzed independently and without across-cohort comparisons.

In the Adu-Oppong cohort, nine *Escherichia* strains were enriched in patients with UTI compared to controls, while two strains MAG398 (*L. jensenii A*) and MAG562 (*S. aureus*) were significantly more abundant in controls (**Fig. 5a** and **Table S2)**. In the comparison between controls and relapse UTIs in the Neugent cohort, 14 *Escherichia* strains were significantly enriched in relapse UTIs, while six strains (e.g., *W. sp002849225*, *S. haemolyticus*, *S. hominis*, and *Corynebacterium pyruviciproducens*) were significantly more abundant in control samples (**Fig. 5b** and **Table S3).** When comparing remission with the relapse UTIs, 13 *Escherichia* strains were enriched in relapse UTIs, while six strains from *W. sp002849225*, *S. haemolyticus*, *Prevotella bivia*, *S. epidermidis,* and *S. hominis* were significantly more abundant in remission samples (**Fig. 5c** and **Table S4).** No differentially abundant strains were identified between controls and remission UTIs.

**Figure 5.**
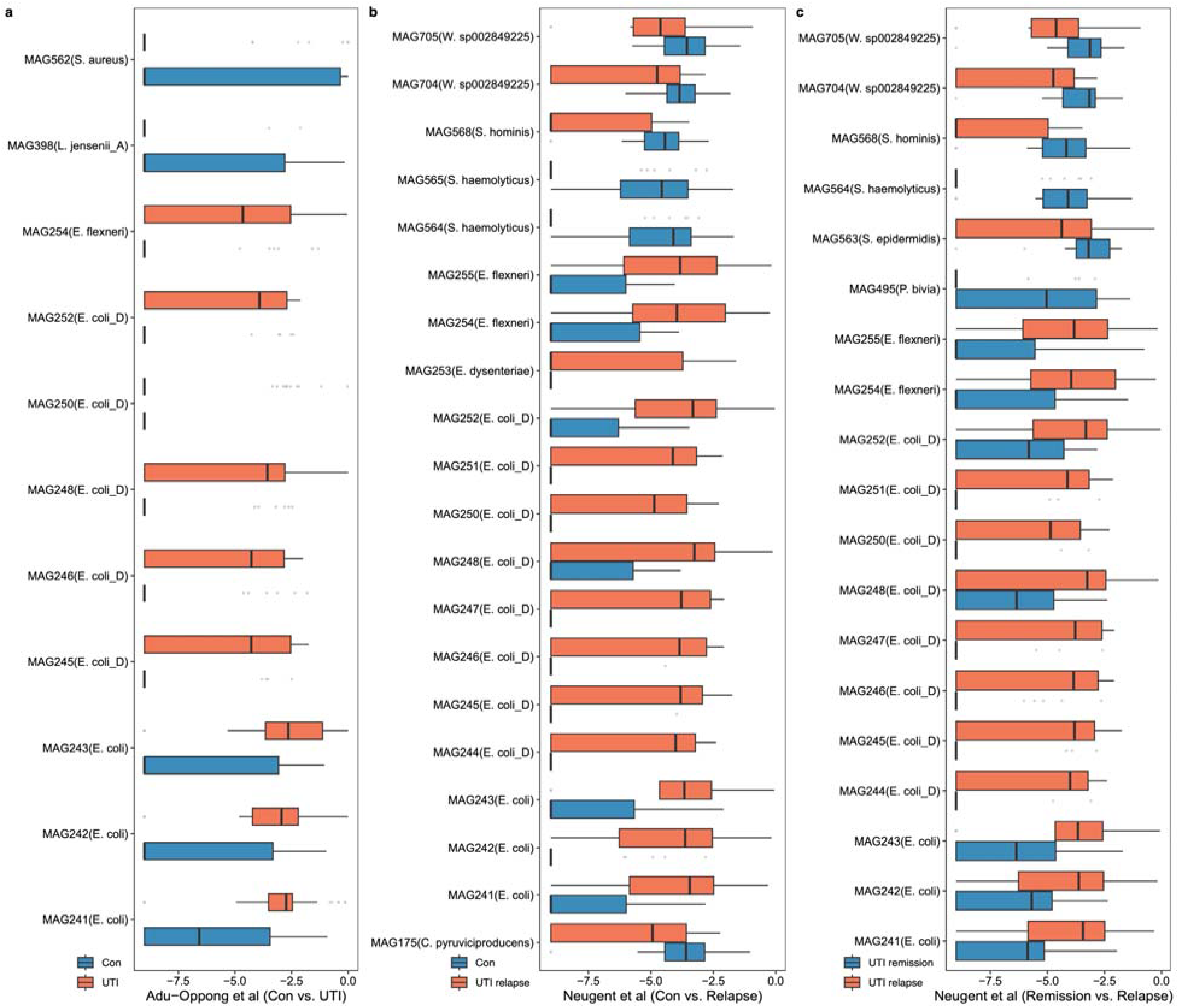
Identification of UTI-associated urinary strains. **a**, The abundance distribution of differential abundant nrMAGs identified in controls vs. patients with UTI in Adu-Oppong. **b**, The abundance distribution of differential abundant nrMAGs identified in No-UTI history vs. recurrent UTI relapse patients in the Neugent cohort. **c**, The abundance distribution of differential abundant nrMAGs identified in remission UTI vs. recurrent UTI relapse patients in the Neugent cohort. Differential abundant nrMAGs were identified using ANCOM, with a Benjamini–Hochberg correction at a 5% level of significance, adjusted for age and sex when available.

Among the three comparisons (controls vs. UTI in the Adu-Oppong cohort, controls vs. relapse, and remission vs. relapse in the Neugent cohort), nine overlapping differentially abundant nrMAGs were identified (**Fig. 5 and Table S2-4**). Of these overlapping nrMAGs, all of them were *Escherichia* strains. Collectively, these metagenomic analysis-based findings support *Escherichia coli* as the primary etiologic agent associated with UTIs, which is consistent with traditional culture-based methods^49^. Also, phylogenetic analysis of *E. coli* genomes further revealed that *E. coli* identified from non-UTI samples were interspersed with those from UTI samples across the phylogenetic tree, without forming a distinct lineage-specific cluster (**Fig. S8**). Together, these findings suggest that the transition from urinary colonization to infection may associated with expansion of *E. coli* populations rather than acquisition of a genetically distinct lineage.

### Mining AMPs from the urinary microbes via deep learning

Bacteria interact with each other through diverse mechanisms, including the production and release of AMPs^50^. Within microbial communities, AMPs can function as selective inhibitory molecules that shape composition of microbial communities by suppressing competing taxa^51^. Beyond their ecological role, AMPs represent a promising class of next-generation antimicrobial agents^52^ because of their potent activity, unique mechanisms of action, and reduced susceptibility to conventional antibiotic resistance^53,54^. However, large-scale discovery of novel AMPs through experimental screening remains time-consuming, labor-intensive, and costly^55^. Recent advances in machine learning have provided powerful approaches to accelerate antimicrobial discovery^56^, including AMP identification from complex microbiome datasets^57^. Given the urgent need for alternative UTI therapeutics in the era of increasing antibiotic resistance, we investigated whether urinary microbiome-encoded AMPs could be identified as potential inhibitors of uropathogens like *E. coli*.

To mine for genes encoding potential AMPs, we first predicted small open-reading frames (sORFs; 5–50 amino acids in length) from the 705 nrMAG genomes. A total of 74,771,567 sORFs were predicted (**Fig. 1b**). Given that computationally predicted sORFs may not be translated into peptides, we mapped these sORFs to 570,420 known proteins in the Swiss-Prot^58^ database to identify likely expressed sORFs. After filtering, 14,201 likely expressed sORFs were retained. We then applied four established machine learning-based AMP prediction tools, including AI4AMP^59^, AMPlify^60^, AMPScanner v2^61^, and iAMPCN^62^ to predict AMPs. Only sORFs predicted as AMPs by all four tools were retained, resulting in 1,569 high-confidence putative AMPs. However, these tools predict general AMP properties rather than activity specifically against *E. coli*. Moreover, experimental validation of 1,569 candidates in the laboratory would be impractical. Therefore, we developed four complementary predictive models (Random Forest: RF, Support Vector Machine: SVM, Long Short-Term Memory: LSTM, and Convolutional Neural Network: CNN) to prioritize the most promising AMP candidates targeting *E. coli* (**Fig. 6a**).

**Figure 6.**
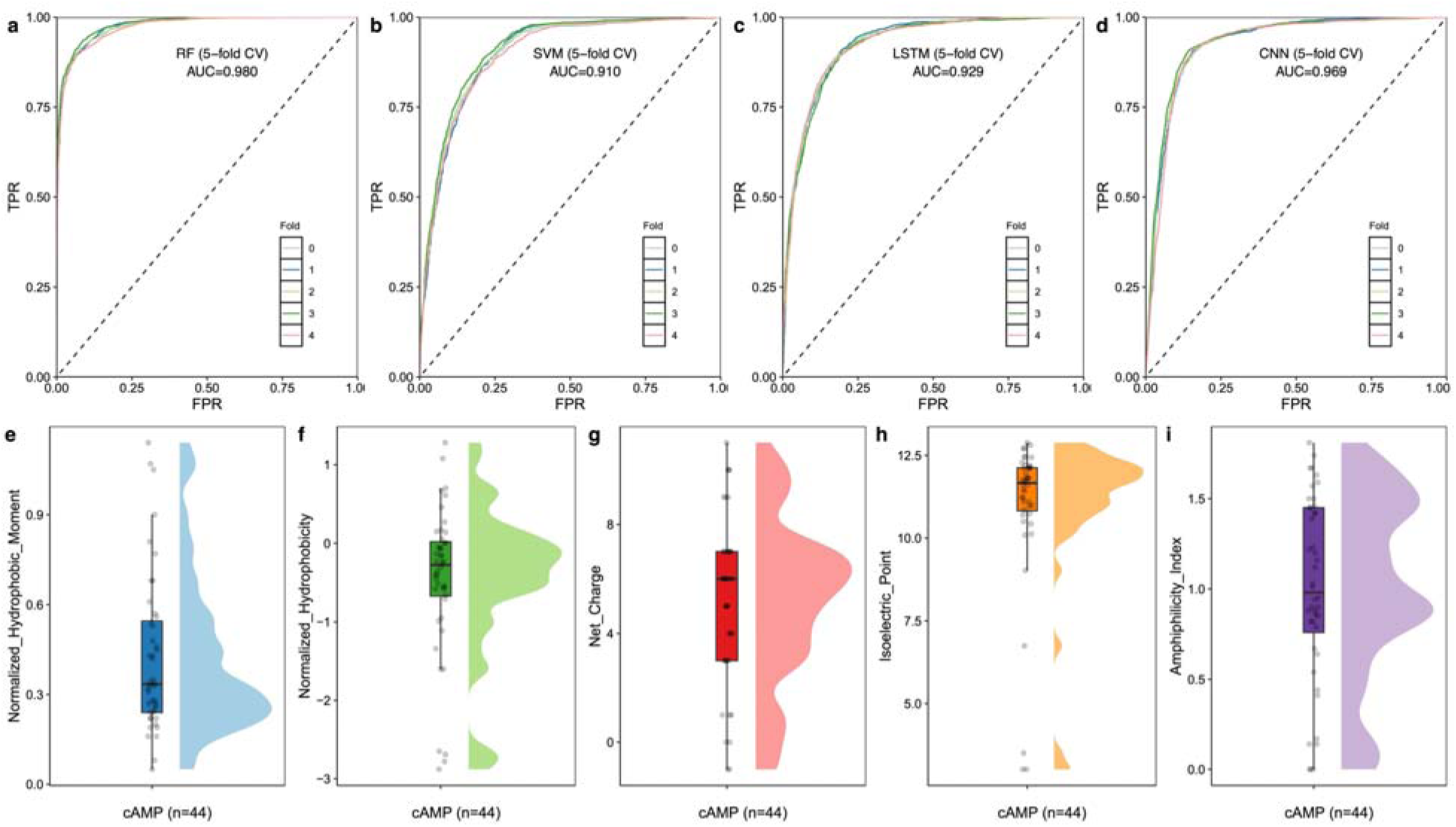
Identification of candidate AMPs from HUMC genomes. **a.** Schematic of the computational approach for the discovery of AMPs from HUMC. The AUROC curves of Random Forest (**b**), SVM (**c**) LSTM (**d**), CNN (**e**), trained with AMPs and non-AMPs with 5-fold cross-validation. Physicochemical features of 44 candidate AMPs on normalized hydrophobic moment (**f**), normalized hydrophobicity (**g**), net charge (**h**), Isoelectric point (**i**), and amphiphilicity index (**j**).

To train the RF, SVM, LSTM, and CNN models, we collected 11,320 AMPs known to target *E. coli* from dbAMP^63^, DRAMP^64^, and DBAASP^65^, along with 11,320 non-AMP sequence from the UniProt^66^ database (**Fig. 1b**). All four models were trained with both AMPs and non-AMPs using 5-fold cross-validation, achieving high classification performance with AUROC values of 0.98 (RF), 0.91 (SVM), 0.93 (LSTM), and 0.97 (CNN) (**Fig. 6a-d**), indicating high predictive performance for identifying AMPs with potential activity against *E. coli*.

We then employed these models to prioritize anti-*E. coli* AMPs from the 1,569 high potential AMPs. Only candidates predicted to exhibit anti-*E. coli* activity by all four models were retained, resulting in 44 candidate AMPs. The physicochemical features of the 44 candidate AMPs were calculated using the Database of Antimicrobial Activity and Structure of Peptides (DBAASP)^65^ server, including normalized hydrophobic moment, normalized hydrophobicity, net charge, isoelectric point, and amphiphilicity index (**Fig. 6e-i, Table S5)**. These candidate AMPs exhibited diverse physicochemical properties, with net charges ranging from -1 to +11, isoelectric points between 3.01 and 12.88, and varying levels of hydrophobicity (−2.88∼1.28) and amphiphilicity (0∼1.81).

### Laboratory testing of candidate AMPs targeting uropathogenic *E. coli* and other pathogens

To assess the activity of the 44 candidate AMPs against uropathogenic *E. coli* (UPEC), we chemically synthesized these peptides. Five peptides were insufficiently soluble in PBS for downstream assays, whereas the remaining 39 soluble candidate AMPs were assessed for antimicrobial activity. Minimal inhibitory concentration (MIC) assays against UPEC strain CFT073 revealed that AMPs 14 and 21 exhibited the strongest antimicrobial activity (**Fig. 7a**). To gain insight into the structural features underlying the observed antimicrobial activity, we predicted the 3D structure of these two lead candidates (**Fig. 7b**) using AlphaFold 3^67^. Secondary structure assignment of the predicted models indicated predominantly α-helical conformations for both peptides, a structural feature associated with antimicrobial activity through membrane-interactive molecular surface signatures^68^.

**Figure 7.**
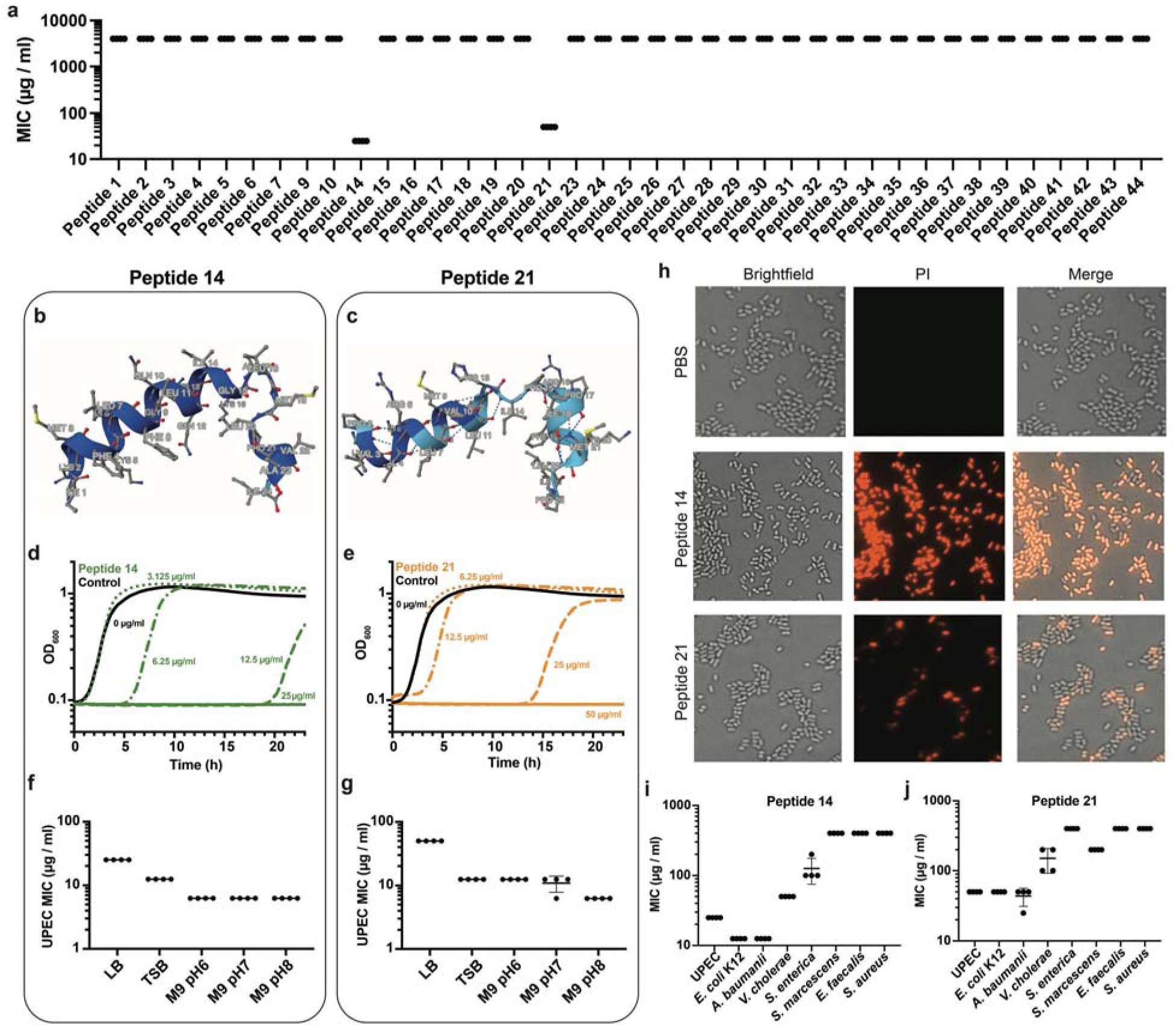
*In vitro* testing of candidate AMPs against uropathogenic and common pathogens. **a.** Minimal inhibitory concentrations (MIC) values of all 44 tested peptides against UPEC in LB (n = 4). 3D structures of peptides 14 (**b**) and 21(**c**), predicted by Alpha-fold 3. Growth curves of UPEC (measured by optical density at 600 nm; OD_600_) over 24 hours in the presence of varying concentrations of peptide 14 (**d**) and 21(**e**). Stability of antimicrobial activity for peptide 14 (f) and 21 (g) against UPEC was assessed across different media types (LB, TSB) and pH ranges (M9 at pH 6, 7, and 8). **h**, Phase-contrast and fluorescence microscopy images as well as a merged image of propidium iodide (PI)–stained UPEC cells treated with 400µg/ml of peptide 14 and 21 as well as PBS. MIC values for peptide 14 (**I**) and 21 (**J**) against *Vibrio cholerae* (contemporary strain from the 2010 outbreak in Haiti)*, E. coli* K12 (DH5α)*, Salmonella enterica* serovar Typimurium (SL1344)*, Serratia marcescens* (ATCC 13880)*, Acinetobacter baumannii* (ATCC 17978)*, Staphylococcus aureus* HG003*, and Enterococcus faecalis* (V583) in LB.

Assays of growth kinetics demonstrated that both peptides inhibited UPEC proliferation in a dose-dependent manner, with AMPs 14 and 21 achieving near-complete growth suppression at concentrations of 25 µg/ml and 50 µg/ml, respectively (**Fig. 7b**). Both peptides also inhibited UPEC growth in a concentration dependent manner under diverse conditions, including rich media (LB and TSB) and minimal media at pH 6, 7 and 8, reflecting the stability of their anti-UPEC activity in diverse conditions (**Fig. 7 f-g**). Most AMPs lead to cell death that can be identified with propidium iodide staining, which detects membrane depolarization. Consistent with this mechanism, microscopy showed that by 15 minutes after treatment with either AMP 14 or 21 there was strong PI staining of UPEC (**Fig. 7h**).

Finally, we investigated the spectrum of activity of these two peptides against a wide variety of Gram-negative and Gram-positive organisms. We found that both peptide 14 and peptide 21 inhibited growth of most of the Gram-negative bacteria tested but not the Gram-positive isolates (**Fig. 7i-j**). Both peptides showed high activity against *E. coli K12, V. cholerae,* and *A. baumanii* in addition to UPEC. There were differences in the specificities of the two peptides, with peptide 14 showing some activity against *Salmonella enterica* and peptide 21 exhibiting activity against *Serratia marcescens*. Thus, these two peptides, derived from the genome sequences of *Finegoldia magna_H* and *Corynebacterium sp001807205*, respectively, may be useful in the treatment of UTIs caused by UPEC as well as infections caused by other Gram-negative bacteria.

## DISCUSSION

Here, we present an extensive human urinary microbial gene and genome catalog generated from whole-metagenome shotgun sequencing data across four publicly available cohorts. This catalog comprises nearly 1.3 million non-redundant microbial genes and 705 non-redundant MAGs, substantially expanding the genomic landscape of the human urinary microbiome. This catalog provides a unique resource to deepen our understanding of the human urinary microbiome and its role in UTIs, serving as a valuable reference source for future human urinary microbiome studies. Further, construction of this microbial genome catalog, enabled a new strain-level perspective for understanding the human microbiome and UTIs and facilitated discovery of new AMPs for UTI therapeutics.

In non-UTI urinary samples, microbial taxa such as *Bifidobacterium longum*, and *Lactobacillus iners*, and *Prevotella colorans* were detected, further supporting the notion that the urinary tract harbors a considerable resident microbial community even in the absence of clinically diagnosed UTI^13^. The microbial diversity and composition in the urinary microbiome were different across different cohorts, potentially due to inherent differences between cohorts, such as host genetics, diet, menopausal status, and age, all of which have previously been associated with urinary microbiome^2^.

Importantly, several common pathogenic bacteria, including *E. coli*, *K. pneumoniae*, and *S. aureus,* were also detected in some non-UTI individuals. However, differential abundance analysis showed that these species were significantly enriched in patients with UTI, suggesting that they are generally present at substantially lower abundances in individuals without infection. This observation is intriguing, as it suggests that host defense mechanisms and/or interactions within the resident urinary microbiota may restrict the overgrowth of uropathogens, thereby preventing the development of symptomatic UTI. Notably, comparative genomic analysis of *E. coli* strains from UTI and non-UTI individuals did not reveal substantial genetic differences, suggesting that the presence of potentially uropathogenic *E. coli* alone may not be sufficient to determine infection outcomes. Instead, the transition from colonization to symptomatic infection may be influenced by multiple factors, including bacterial abundance, microbial community context, and host susceptibility^69^. These findings also highlight an important limitation of conventional urine culture: although urine culture remains essential for clinical diagnosis and antimicrobial susceptibility testing, it provides limited information about the broader urinary microbial ecosystem and may fail to capture fastidious, low-abundance, or difficult-to-cultivate organisms^70^. In some cases, the presence of *E. coli* may represent asymptomatic carriage or low-level colonization rather than an active infection requiring treatment. Conversely, culture-based approaches may not fully distinguish microbial colonization from infection when bacterial detection occurs in the context of complex urinary microbial communities. Therefore, microbial detection alone should be interpreted together with microbial abundance, strain-level characteristics, and host clinical context to better distinguish true infection from colonization.

Given the extensive use of antibiotics for UTI treatment, characterizing antimicrobial resistance potential within urinary microbial communities is critical. Comparative analyses across two independent case-control cohorts demonstrated that urinary microbiomes from patients with UTI exhibited enrichment of specific ARGs and VF genes compared with non-UTI controls, indicating functional differences associated with infection status. These findings are consistent with previous studies based on bacterial isolation and culture^32,71–73^. The enriched ARGs in UTI-associated microbiomes were linked to resistance mechanisms against multiple antibiotic classes, including fluoroquinolones, carbapenems, macrolides, and tetracyclines. Our gene catalog offers a unique opportunity to characterize the genomic potential for antibiotic resistance beyond extensively studied bacterial isolates, including microbial populations that may be difficult to recover using conventional culture-based approaches. In addition, *Escherichia* strains were consistently enriched in UTI samples across independent cohorts, highlighting their important role in UTI. This is consistent with the fact that UPEC is considered the most common cause of both uncomplicated and complicated UTIs^74^. In future studies, it will be valuable to further analyze whether some of the *E. coli* identified here differ from traditional UPEC strains in their virulence gene repertoire.

Microbial communities, which are metabolically diverse^30^, produce many antimicrobial factors including AMPs^75^. AMPs are considered potential next-generation antimicrobial agents to combat increasingly resistant pathogens^54^. Indeed, several AMPs, including cathelicidin^76,77^, hBD1^78^, and RNase7^79^, have been shown to have potential against urinary pathogens. Our 705 nr MAGs generated through d*e novo* assembly and binning offered a valuable resource to extract potential AMPs. We first leveraged an ensemble strategy by integrating predictions from four independent machine learning models to identify candidate AMPs through a consensus voting approach, thereby improving robustness and reducing model-specific bias. We then developed a tailored machine learning model to further refine these candidates, specifically prioritizing peptides with predicted activity against *E. coli*. Through this two-stage framework, we narrowed down a long list of candidates to 44 potential AMPs for laboratory testing. We established that two of the candidate AMPs showed antimicrobial activity against UPEC in various media types. Structural prediction further suggested that these peptides may exert antimicrobial activity through mechanisms consistent with previously characterized AMPs, including membrane disruption and depolarization of Gram-negative bacteria. Our results suggest that urinary microbial genomes represent an underexplored reservoir for the discovery of novel antimicrobial molecules.

Several limitations should be noted in this study. First, most of the urinary samples were clean-catch midstream urine. Therefore, they may represent a mixed microbial community with bacteria from the bladder, periurethra or the genital tract. Second, even though we adjusted for potential confounders in our statistical models, we were unable to assess some covariates such as medication, diet, and psychological stress that are not publicly available. Third, a large proportion of unmapped sORFs were discarded, yet these could represent novel peptides lacking homology in existing protein databases. Finally, animal experiments are needed to assess the efficacy of candidate AMPs in model UTIs, their toxicity to human cells, and their bioavailability, as well as to translate these findings into clinical applications. Despite these limitations, the high-quality gene and genome catalogs presented here will enhance urinary microbiome research, uncovering previously uncharted genomic information that can advance our understanding of the role of urinary microbes in UTIs.

In conclusion, we created an extensive human urinary microbial gene and genome catalog that will serve as a valuable resource for investigating the role of the urinary microbiome in health and disease. Our findings show that UTI samples exhibit increased abundance of ARGs and VF genes, reflecting enrichment of genomic features associated with pathogenic potential during infection. We further observed that *E. coli* is present in both non-UTI and UTI individuals but is significantly more abundant in patients with UTI, indicating that disease is associated with microbial overgrowth rather than exclusive pathogen presence. We further developed an AI-based pipeline to identify potential AMPs active against uropathogenic *E. coli*. The success of this approach suggests that specific urinary microbes have the potential to inhibit the growth of uropathogenic *E. coli* via the release of AMPs. Together, our study establishes a foundational resource for the urinary microbiome and highlights its potential to inform mechanistic understanding of UTI pathogenesis and the development of microbiome-based therapeutic strategies.

## METHODS

### Study cohorts

We systematically identified urinary metagenomic sequencing studies from keyword searches in PubMed and online repositories (i.e., NCBI and ENA). We included samples with publicly available raw shotgun metagenomic sequencing data (paired fastq files) and metadata. All the sequencing data were downloaded from online repositories or links provided in the original publications. We did not include any studies that required additional ethics committee approvals or authorizations for access. A total of 450 urinary microbiome samples from four independent cohorts were analyzed in this study (**Fig. 2a**). A total of 177 (non-UTIs), 149 (asymptomatic, contaminated, insignificant, no growth, and positive), 49 (unknown), and 75 (non-UTIs, UTI remission, and UTI relapse) urinary microbiome were included from Adebayo, Adu-Oppong, Moustafa, and Neugent cohorts, respectively. Specifically, only symptomatic patients in the Adu-Oppong cohort confirmed by *in vitro* positive culture (specimen had significantly growth of one or two uropathogens) were included in the case-control comparison.

### Metagenome quality control and assembly

The shotgun metagenomic sequences were preprocessed and quality checked using the modules available at MetaWRAP (v1.3.2)^80^, including trimming the raw sequence reads and removing human contamination for each of the sequenced samples. Next, the clean reads from the sequencing samples were assembled with the metaWRAP-Assembly module using metaSPAdes (v3.13.0)^81^.

### Microbial gene catalog and gene annotation

A total of 201,051,463 assembled contigs were used for gene prediction by Prodigal (v2.6.3)^33^. After removing the incomplete genes, those with a start and stop codon were retained for further analyses. All complete genes were clustered at the protein level following UniRef guidelines at 100%^82^. Gene abundance was estimated using Salmon (v0.13.1)^83^. We then annotated the genes from the HUMC using eggNOG-mapper (v2.1.7)^36^ with a broad range of databases, including COGs^84^ (Clusters of Orthologous Genes), KEGG^85^ (Kyoto Encyclopedia of Genes and Genomes), PFAMs^86^ (protein families), and ECs^87^ (Enzyme Commission). Additionally, we annotated antibiotic resistance genes and virulence factors using CARD^42^ (Comprehensive Antibiotic Resistance Database) and VFDB^45^ (Virulence Factor Database), respectively.

### Construction of microbial genomes

The assembled contigs were binned into bins using two metagenomic binning tools: MetaWRAP (v1.3.2)^80^ [including metaBAT2^88^ (v2.12.1), MaxBin2^89^ (v2.2.6), CONCOCT^90^ (v1.0.0)] and VAMB^91^ (v3.0.2). MAGs refinement was performed using the bin_refinement module of metaWRAP. CheckM^92^ (v1.0.12) was used to estimate the completeness and contamination of the bins, with a minimum completeness threshold of 50% and a maximum contamination threshold of 10%.

### De-replication of MAGs and genome annotation

All 2407 MAGs were de-replicated into non-redundant MAGs (nrMAGs) using dRep (v3.0.0)^93^ (≥50% genome completeness and ≤5% contamination)^50^. Initially, MAGs were divided into primary clusters using Mash at a 90% Mash ANI^94^. Next, each primary cluster was used to form secondary clusters at the threshold of 99% ANI with at least 30% overlap between genomes. Taxonomic annotation of all nrMAGs was conducted using GTDB-Tk^95^ (v.1.4.1) based on the Genome Taxonomy Database (http://gtdb.ecogenomic.org/)^46^. The metaWRAP-Quant_bins module, integrated with Salmon (v0.13.1)^83^, was used to estimate the abundance of each nrMAG in the metagenomic samples.

### Prediction of sORFs from bacteria genomes

To explore the hypothesis that bacteria may interact with uropathogen *E. coli* via the release of antimicrobial peptide, we first predict sORFs from seven UTI-related bacteria genomes using the ‘getorf’ function of the EMBOSS^96^ software package (version 6.6.0). To ensure that the sORFs predicted in silico are likely-expressed to proteins/peptides, we examined them with known proteins in Swiss-Prot^58^, an annotated protein sequence database. When an sORF can match a known protein, it suggests that the sORF is not a random sequence, but rather a potential protein or peptide that may function in a similar biological context. This matching provides strong evidence that this sORF is likely to be expressed as a functional protein or peptide.

### Identification of candidate AMPs from likely expressed sORFs using deep learning models

To identify potential AMPs from the likely expressed sORFs, we applied four existing computational tools, including AI4AMP^59^, AMPlify^60^, AMPScanner v2^61^, and iAMPCN^62^ to predict AMPs. These tools identify peptides with general antimicrobial potential but do not predict activity against specific bacterial species. Therefore, they do not indicate whether the predicted peptides can inhibit *E. coli*. To address this limitation, we developed and trained our own models to predict *E. coli* inhibitory activity. To train the machine learning models, we first collected AMPs from five databases (dbAMP^63^, DRAMP^64^, APD3^97^, SATPdb^98^, and DBAASP^65^). Among these five databases, only three (dbAMP, DRAMP, and DBAASP) provide information on target bacteria of given AMPs. A total of 11,320 AMPs that can target *E. coli* were selected. Additionally, 11,320 Non-AMPs from the UniProt^66^ database by setting the ‘subcellular location’ filter to the cytoplasm and removing any entry that matched the following keywords: antimicrobial, antibiotic, antiviral, antifungal, effector or excreted (downloaded as of 10 May 2023)^99^. AMPs and Non-AMPs that were longer than 100 AAs or shorter than 2 AAs were excluded. The AMP and Non-AMPs were used to train the RF, SVM, LSTM, and CNN models. The training data are preprocessed, split into training (4-folds) and testing (1-fold) sets, and transformed into a tensor format for compatibility with the models. The performance of the models was measured by AUROC (area under the receiver operating characteristic curve). The non-redundant and likely expressed sORFs sequences from HUMC strains were converted into a numeric format to feed two pre-trained deep learning models. Only sORFs predicted as AMP by both LSTM and CNN models were considered as a strong candidate AMP.

### Peptide synthesis

All 44 candidate AMPs used in this study were synthesized by AAPPTec (KY, USA). The purity of all AMPs was determined by high-performance liquid chromatography, and all had a purity greater than 98%.

### Minimum inhibitory concentration determination

All 44 peptides were dissolved in sterile-filtered 1× PBS to a concentration of 20 mg/ml. Peptides 8, 11, 12, 13 and 22 did not fully dissolve and were not used for further studies. 1:10 serial dilutions of AMPs were added to sterile 96-well plates with LB (lysogeny broth) or the respective growth medium indicated.

All bacterial strains were diluted from an overnight grown culture to a final concentration of 1:5000 into 96-well plates with different AMP solutions diluted serially. After incubating at 37°C for 20 h, the MIC was determined as the minimum concentration of AMPs where bacteria showed no detectable growth. Growth was determined as a OD_600_ ≥ 50% of the positive control, containing no AMPs. The OD_600_ of each well was measured on a microplate reader. All experiments were performed with four independent replicates.

### Growth kinetics

Growth kinetics were performed in transparent 96-well plates (Greiner) with 150 µl culture volume. UPEC bacteria were grown in a pre-culture for ∼16 h in LB with aeration and shaking at 37°C. For growth assays, pre-cultures were diluted 1:10000 in LB with or without different concentrations of peptide 14 or 21. The OD_600_ was monitored every 10 min at 37°C with shaking. For presentation of data, the mean of four independent growth curves was plotted.

### Microscopy assay

100 ul UPEC strain from the exponential phase were diluted 1000-fold with sterile PBS and then centrifuged for two minutes (3000 rpm). After centrifugation, the supernatant was discarded. The precipitate was well mixed with AMPs or controls and then transferred onto 1.5% agarose pads. Propidium Iodide was added directly into the melted agarose at 55 °C immediately before preparing the pads.

### Statistical analysis

The comparison between the genes from the HUMC and HRGM catalogs was calculated by cd-hit. The difference in gene prevalence between control and patients with UTI was measured by chi-squared test with the Benjamini–Hochberg correction implemented in the p.adjust function in R. Differential abundant ARG or VF genes were identified using Wilcoxon–Mann–Whitney test, P values were corrected using the Benjamini–Hochberg procedure. The phylogenetic tree of nrMAGs was built using PhyloPhlAn^100^ (v3.0.58) and then visualized using iTOL^101^ (https://itol.embl.de/). nrMAGs annotated to such as *Delftia, Stenotrophomonas, Ralstonia,* and *Bradyrizobium* were exluded as they were likely contamination. Microbial richness was calculated at the nrMAG level using the ‘vegan’ R package. Differential abundant nrMAGs were identified using ANCOM^48^, with a Benjamini–Hochberg correction at a 5% level of significance, adjusted for age and sex when available. The three-dimensional structures of the candidate antimicrobial peptides were predicted using AlphaFold3^67^. All statistical analyses were performed in R (version 4.4.0) or Python (version 3.10.7).

### Data availability

The metagenomic sequencing data in this study can be downloaded via NCBI (BioProject ID: PRJNA385350; PRJNA700071; PRJEB36610; PRJNA801448). HUMC is available on Figshare (https://figshare.com/). Training data of our deep learning models are publicly available at: dbAMP (http://awi.cuhk.edu.cn/dbAMP), DRAMP (http://dramp.cpu-bioinfor.org/), DBAASP (https://dbaasp.org/), and Uniport (https://www.uniprot.org/).

### Code availability

The code for the construction of the HUMC and statistical analysis and visualization is available in the GitHub repository (https://github.com/KelabatOSU/UTI).

## Acknowledgements

S.K. acknowledges funding support from the Department of Internal Medicine pilot grant, OSUCCC Molecular Carcinogenesis and Chemoprevention Program pilot grant at The Ohio State University, and the Women’s Health Interdisciplinary Stress Program of Research (WHISPR) Pilot Grant from the Connors Center at Brigham and Women’s Hospital. Y.-Y.L. acknowledges funding support from NIH (R01AI141529). X.-W.W. acknowledges funding support from NIH (K25HL166208). F.G.Z. and M.K.W. are supported by NIH (R01AI042347) and HHMI. The research by F.G.Z. was supported by the Life Sciences Research Foundation (Zingl-2024HHMI).

## Author contributions

S.K. conceived the project. S.K. and Y.-Y.L. designed the project. S.K. analyzed all the human urinary microbiome data. S.K. applied machine learning models in AMP mining, with assistance from X.-W.W., M.K.W., and Y.-Y.L. interpreted the results of the human data analysis. S.K. and F.G.Z conducted *in vitro* experiments. M.K.W. interpreted results. S.K. drafted the manuscript. Y.-Y.L. X.-W.W., M.K.W., F.G.Z., and S.T.W. edited the manuscript. All authors reviewed and approved the manuscript.

## Competing interests

The authors declare no competing interests.

## Notes

https://github.com/KelabatOSU/UTI

### Competing Interest Statement

The authors have declared no competing interest.

